# Inhibition of gibberellin accumulation by water deficiency promotes fast and long-term ‘drought avoidance’ responses in tomato

**DOI:** 10.1101/2021.04.13.439675

**Authors:** Hagai Shohat, Hadar Cheriker, Himabindu Vasuki, Natanella Illouz-Eliaz, Shula Blum, Ziva Amsellem, Danuše Tarkowská, Asaph Aharoni, Yuval Eshed, David Weiss

## Abstract

Plants reduce transpiration to avoid dehydration during drought episodes by stomatal closure and inhibition of canopy growth. While abscisic acid (ABA) has a primary role in ‘drought avoidance’, previous studies suggest that gibberellin (GA), might also be involved. Here we show in tomato (*Solanum lycopersicum*) that shortage of water inhibited the expression of the GA biosynthesis genes *GA20 oxidas*e*1* (*GA20ox1*) and *GA20ox2* and induced the GA-deactivating gene *GA2ox7* in leaves and guard cells, resulting in reduced bioactive GA levels. Drought regulation of GA metabolism was mediated by ABA-dependent and independent pathways, and by the transcription factor *DEHYDRATION RESPONSIVE ELEMENT BINDING (DREB), TINY1*. Mutations in *GA20ox1* and *GA20ox2* reduced water loss due to the smaller canopy area. On the other hand, loss of *GA2ox7* did not affect leaf size, but attenuated stomatal response to water deficiency; during soil dehydration, *ga2ox7* plants closed their stomata and reduced transpiration later than WT, suggesting that *ga2ox7* stomata are hyposensitive to soil dehydration. Together, the results suggest that drought-induced GA deactivation in guard cells contributes to stomatal closure at the early stages of soil dehydration, whereas inhibition of GA synthesis in leaves promotes mainly the long-term reduction in canopy growth to reduce transpiration area.

## INTRODUCTION

Drought is a common and devastating abiotic stress that reduces crop yield worldwide (Fahad et al., 2017). Water deficiency inhibits plant growth, flowering and fruit development (Gupta et al., 2020). It also suppresses directly and indirectly major biochemical pathways, including photosynthesis and primary carbon metabolism (Tardieu et al., 2018). Plants use three major strategies to cope with and/or adapt to drought: drought escape, drought avoidance and drought tolerance (Kooyers, 2015). To escape from water-deficit stress, plants complete their life cycle before drought becomes severe. Tolerance to drought is acquired by osmotic adjustment, ROS scavenging and activation of stress-related genes. Plants can also avoid dehydration during transient periods of water deficiency by stomatal closure and growth suppression, both leading to reduced transpiration. Drought avoidance is regulated primarily by the stress-hormone abscisic acid (ABA) (Cutler et al., 2010). However, several studies suggest that changes in the levels of the growth-promoting hormone, gibberellin (GA), may also be involved (Colebrook et al., 2014).

GA promotes major developmental processes throughout the plant life cycle, including seed germination, shoot elongation, leaf expansion, flowering and fruit development (Yamaguchi, 2008). The nuclear proteins DELLA suppress all GA responses by interacting with numerous transcription factors (Hauvermale et al., 2012; Locascio et al., 2013). When GA binds to its receptor GIBBERELLIN-INSENSITIVE DWARF1 (GID1), it induces DELLA degradation via the ubiquitin-proteasome pathway, leading to the activation of GA responses (Daviere and Achard, 2013). GA activity is controlled also at the level of hormone biosynthesis and deactivation. Both endogenous and environmental cues regulate the expression of three small gene families acting late in the GA biosynthetic pathway and coding for 2-oxoglutarate-dependent-dioxygenases (2-ODDs). These include the GA 20-oxidases (GA20ox) that cleave C-20 to generate C-19 GAs, GA 3-oxidases (GA3ox) that form the bioactive GAs, GA_1_ and GA_4_ by 3β-hydroxylation and GA 2-oxidases (GA2oxs) that deactivate bioactive GAs or their C-19 and C-20 precursors (Hedden, 2020). Plants maintain GA homeostasis by feedback response; reduced GA activity upregulates *GA20ox* and *GA3ox* expression and inhibits *GA2ox*, whereas increased GA activity has the opposite effect. These transcriptional feedback and feed-forward regulation is mediated by changes in DELLA levels (Middleton et al., 2012; Fukazawa et al., 2017).

Biotic and abiotic stresses affect GA levels by up-or down-regulating *GA20ox, GA3ox* or *GA2ox* genes (Yamaguchi, 2008). Several studies suggested that water-deficit conditions reduce GA levels (Clebrook et al., 2014); drought induced the expression of *GA2ox* in *Populus* (Zawaski and Busov, 2014), and reduced the levels of bioactive GAs in maize leaves (Nelissen et al., 2018). Moreover, overexpression of drought-related transcription factors from the *DEHYDRATION RESPONSIVE ELEMENT BINDING (DREB)* family in tomato and Arabidopsis reduces GA levels and improves salt and drought tolerance (Li et al., 2012; Magome et al., 2008). Numerous studies demonstrate that low GA activity increases plant tolerance to abiotic stresses, including salt and drought (Achard et al., 2006; Magome et al., 2008; Nir et al., 2017; Illouz-Eliaz et al., 2020). Reduced GA levels lead to the activation of various stress-related genes (Tuna et al., 2008), accumulation of osmolytes (Omena-Garcia et al., 2019) and ROS scavenging enzymes (Achard et al., 2008), all related to drought tolerance. Previously we showed that suppression of GA accumulation in tomato reduced water loss under water-deficit conditions (Nir et al., 2014). Moreover, transgenic tomato plants overexpressing the constitutively active stable tomato DELLA protein *proceraΔ17* (*proΔ17*), exhibited lower whole-plant transpiration due to smaller canopy area and reduced stomatal aperture (Nir et al., 2017). Expressing *proΔ17* specifically in guard cells was sufficient to reduce stomatal aperture without affecting growth, suggesting that this effect of DELLA is cell autonomous. The effect of *proΔ17* on stomatal closure and water loss were suppressed in the ABA-deficient *sitiens* (*sit*) mutant, indicating that these effects of DELLA are ABA-dependent (Nir et al., 2017). High levels of DELLA promoted the expression of the ABA transporter gene *ABA-IMPORTING TRANSPORTER1*.*1* (*AIT1*.*1*) in guard cells, and the *ait1*.*1* mutant suppressed the effect of DELLA on ABA-induced stomatal closure and transpiration (Shohat et al., 2020). These suggest that *AIT1*.*1* mediates, at least partially, the effect of DELLA on stomatal closure. Taken together, these and other results suggest that water deficiency reduces GA levels, which leads to DELLA accumulation. The accumulated DELLA inhibits canopy growth and promotes ABA-induced stomatal closure; both reduce transpiration and water loss. However, a direct link between water availability, changes in GA levels, growth suppression, stomatal closure and lasting avoidance from drought stress was not demonstrated.

To this aim, we have studied how water availability affects GA metabolism in tomato guard cells and leaf tissue and in turn, how this affects transpiration. We show that water deficiency suppressed GA accumulation by downregulating the GA biosynthesis genes *GA20ox1* and *GA20ox2*, and by upregulating the GA deactivating gene *GA2ox7* in both leaf tissue and guard cells. The reduced GA levels had a prolonged impact by inhibiting leaf growth and a short-term impact on stomatal closure. Together, these changes reduced transpiration and promoted ‘drought avoidance’.

## RESULTS

### Water deficiency suppresses GA accumulation in guard cells

A rapid and efficient guard cell isolation procedure (See methods, Shohat et al., 2020) was used to examine if water deficiency affects GA accumulation in guard cells. This rapid procedure was taken to minimize the effect of the isolation process on transcription and hormone metabolism. Tomato M82 (WT) plants were grown under normal irrigation regime, or exposed to water deficiency [15% soil volumetric water content (VWC)] and then guard cells were isolated from leaves number 3 and 4 (top down). Microscopic analysis of the guard-cell enriched samples, stained with neutral red, confirmed the viability of the guard cells, but not of the remaining epidermal cells (Supplemental Fig. 1). We then analyzed GA content in the guard-cells enriched samples and found that water deficiency reduced significantly the levels of the bioactive GAs, GA_1_ and GA_3_, and the C-20 intermediate, GA_12_ (Fig. 1A, Dataset S1). Although GA_3_ is rather rare in plants (Hedden. 2020), previous studies demonstrate its accumulation in tomato (Li et al. 2020). The level of the bioactive GA_4_ was much lower than that of GA_1_ and GA_3_, and was slightly but not significantly higher in the drought treatment.

**Figure 1.**
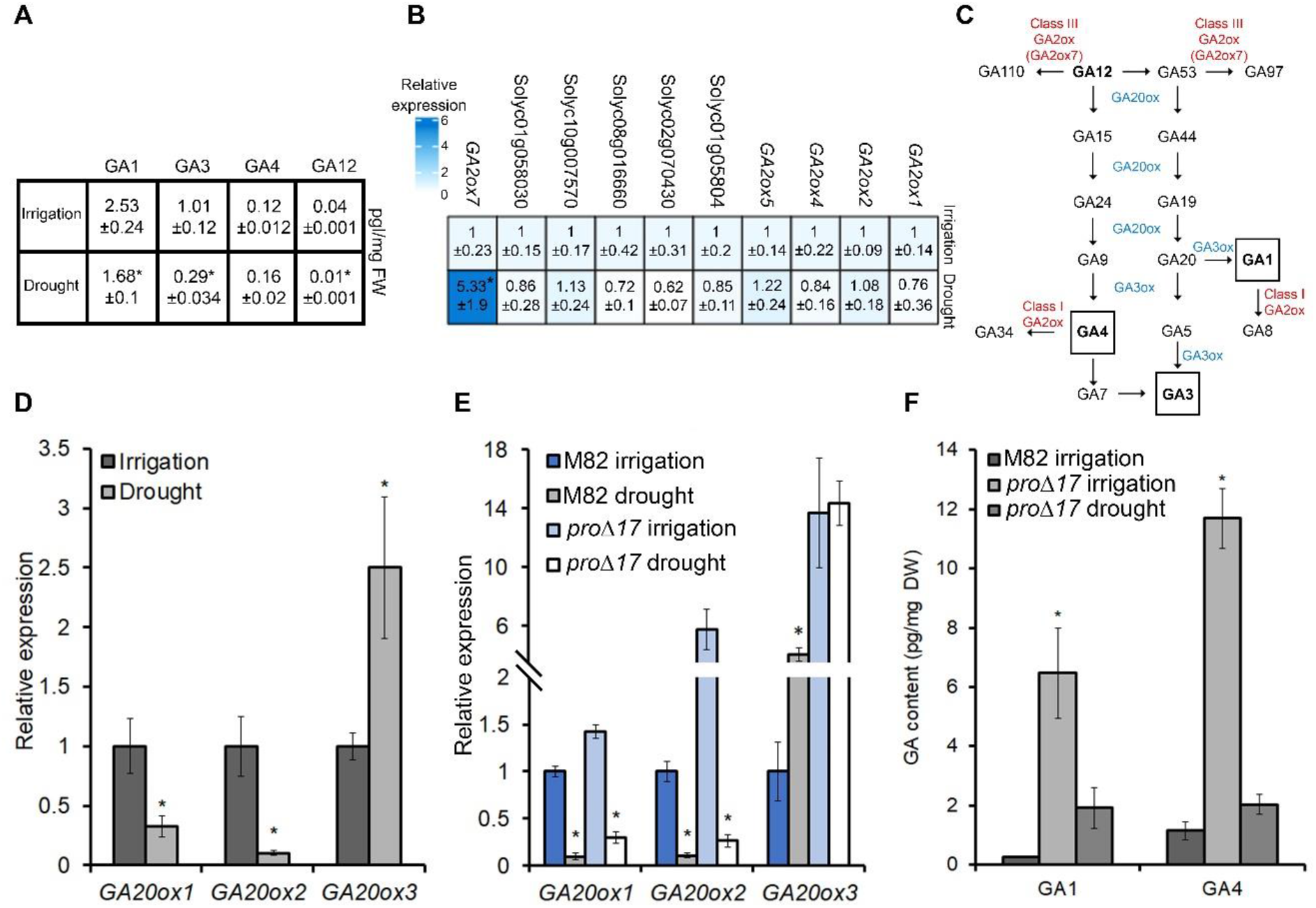
Drought inhibits GA biosynthesis in guard cells and leaf tissue. **(A)** GA levels under irrigation and water-deficit conditions (Soil VWC 15%) in guard cells isolated from leaves number 3 and 4 (top down). Values are means of 4 biological replicates ±SE. FW-fresh weight. **(B)** Heat map showing the relative expression of the tomato *GA2ox* genes in isolated guard cells under irrigation or drought treatments (Soil VWC 15%). Values are means of 4 biological replicates ±SE. **(C)** Scheme of GA metabolism showing the GA biosynthesis enzymes (blue), GA deactivation enzymes (red) and bioactive GAs (black squares). **(D)** *GA20ox1, GA20ox2* and *GA20ox3* expression in isolated guard cells taken from M82 plants that grew with irrigation or exposed to water deficiency (Soil VWC 15%). Values are means of 4 biological replicates ±SE. **(E)** *GA20ox1, GA20ox2* and *GA20ox3* expression in M82 and 35S:*proΔ17* leaves under irrigation or drought conditions (Soil VWC 15%). **(F)** Levels of the bioactive GA_1_ and GA_4_ in leaves of irrigated M82 and 35S:*proΔ17* and drought treated 35S:*proΔ17* (Soil VWC 15%). Values are means of 4 biological replicates ±SE. DW-dry weight. Stars in **A** to **F** represent significant differences between respective treatments by Student’s *t* test (*P < 0*.*05*).

Colebrook et al. (2014) suggest that abiotic stresses reduce GA accumulation by upregulating the GA deactivating gene *GA2ox*. We analyzed the expression of the 11 tomato *GA2ox* genes (Pattison et al., 2015; Chen et al., 2016) in guard-cells enriched samples, following water-deficit treatment (15% soil VWC). Among the 11 *GA2ox* genes, only *GA2ox7* was strongly upregulated by the drought treatment (Fig. 1B). Comparing transcript levels of all *GA2oxs* in isolated WT guard cells using available RNAseq data (Shohat et al., 2020), revealed that *GA2ox7* expression is much higher than all other *GA2oxs* (Supplemental Fig. 2). The strong induction of *GA2ox7* by water deficiency can explain the low level of GA_12_ found under drought conditions, as this C-20 precursor is a substrate of GA2ox7 in tomato (Fig. 1C; Schrager-Lavelle et al., 2019). It is interesting to note that *GA2ox7* is the closest homolog of the Arabidopsis *GA2ox8* (Chen et al., 2016), which is expressed specifically in guard cells (Li et al., 2019).

We next examined whether water deficiency also suppresses GA biosynthesis in guard cells via the inhibition of the GA biosynthesis genes, *GA20ox* or *GA3ox*. Tomato has 8 *GA20ox* and 6 *GA3ox* genes (Pattison et al., 2015). While none of the *GA3ox* genes was downregulated (Supplemental Fig.3), the expression of two *GA20ox* genes, *GA20ox1* and *GA20ox2* was suppressed (Fig. 1D). Surprisingly, the expression of *GA20ox3* was strongly upregulated under water-deficit conditions.

To examine if these changes in GA metabolism are guard-cell-specific, we analyzed the expression of all the above-mentioned genes also in whole-leaf tissue following soil dehydration (15% soil VWC). Similar to guard cells, *GA2ox7* was upregulated and *GA20ox1* and *GA20ox2* were downregulated (Supplemental Fig. 4A-C). The expression of *GA20ox3* was again, strongly upregulated by water-deficit conditions. GA analysis in drought-treated leaves (15% soil VWC) showed strong suppression of GA_1_ but not GA_4_ (Supplemental Fig. 4D, Dataset S1).

### Drought outcompetes the DELLA-induced feedback response to maintain low GA levels under water-deficit conditions

We examined if the drought-induced *GA20ox3* upregulation is a result of a feedback response due to the reduced GA levels (Middleton et al., 2012; Fukazawa et al., 2017). Since the upregulation of GA-biosynthesis genes by reduced GA levels (feedback response) is mediated by the accumulation of DELLA (Middleton et al., 2012), we analyzed the expression of *GA20ox1, GA20ox2* and *GA20ox3* in leaves of WT and transgenic plants over-expressing stable DELLA protein (*35S:proΔ17*, Nir et al., 2017), under irrigation and water deficiency. In well-watered plants, all three genes showed higher expression in *proΔ17* leaves compared to WT (Fig. 1E), suggesting that all are upregulated by the DELLA-mediated feedback response. However, when these plants were exposed to water-deficit conditions, the expression of *GA20ox1* and *GA20ox2*, but not that of *GA20ox3*, was strongly suppressed in both WT and *proΔ17*. We also examined how these changes in gene expression in *proΔ17* affect GA content in leaves. The levels of the two bioactive GAs, GA_1_ and GA_4_, were much higher in *proΔ17* than in WT leaves and strongly reduced under water-deficit conditions (Fig. 1F, Dataset S1,). Moreover, we found a strong reduction in the level of the direct precursor of GA_1,_ GA_20,_ and higher levels of GA_19_, the precursor of GA_20_, indicating an overall inhibition of GA20ox activity (Dataset S1, Fig. 1C; Hedden, 2020). Taken together, these results demonstrate that water deficiency out-competes the effects of DELLA on the transcriptional regulation of *GA20ox1* and *GA20ox2* and keeps them low despite the activation of the feedback response by the accumulated DELLA. This overcomes the mechanism of homeostasis and maintains low GA levels.

### ABA and *DREB-TINY1* regulate GA metabolism under drought conditions

ABA accumulates in tomato leaves under water-deficit conditions (Nir et al., 2017). We examined if ABA mediates the effect of water deficiency on GA metabolism. ABA treatment suppressed the expression of *GA20ox2* and induced the expression of *GA2ox7*, but had no effect on *GA20ox1* (Supplemental Fig. 5A). We further examined the effect of water deficiency on the expression of these three genes in the leaves of the ABA-deficient mutant *sit*. While *GA20ox1* expression was equally downregulated by drought in WT and *sit* (Fig. 2A), the expression of *GA20ox2* was strongly downregulated in WT, but was hardly affected in *sit* (Fig. 2B). The upregulation of *GA2ox7* by water deficiency was partially inhibited in *sit* (Fig. 2C). These results suggest that water-deficit conditions affect GA biosynthesis and deactivation via both ABA-dependent and independent pathways.

**Figure 2.**
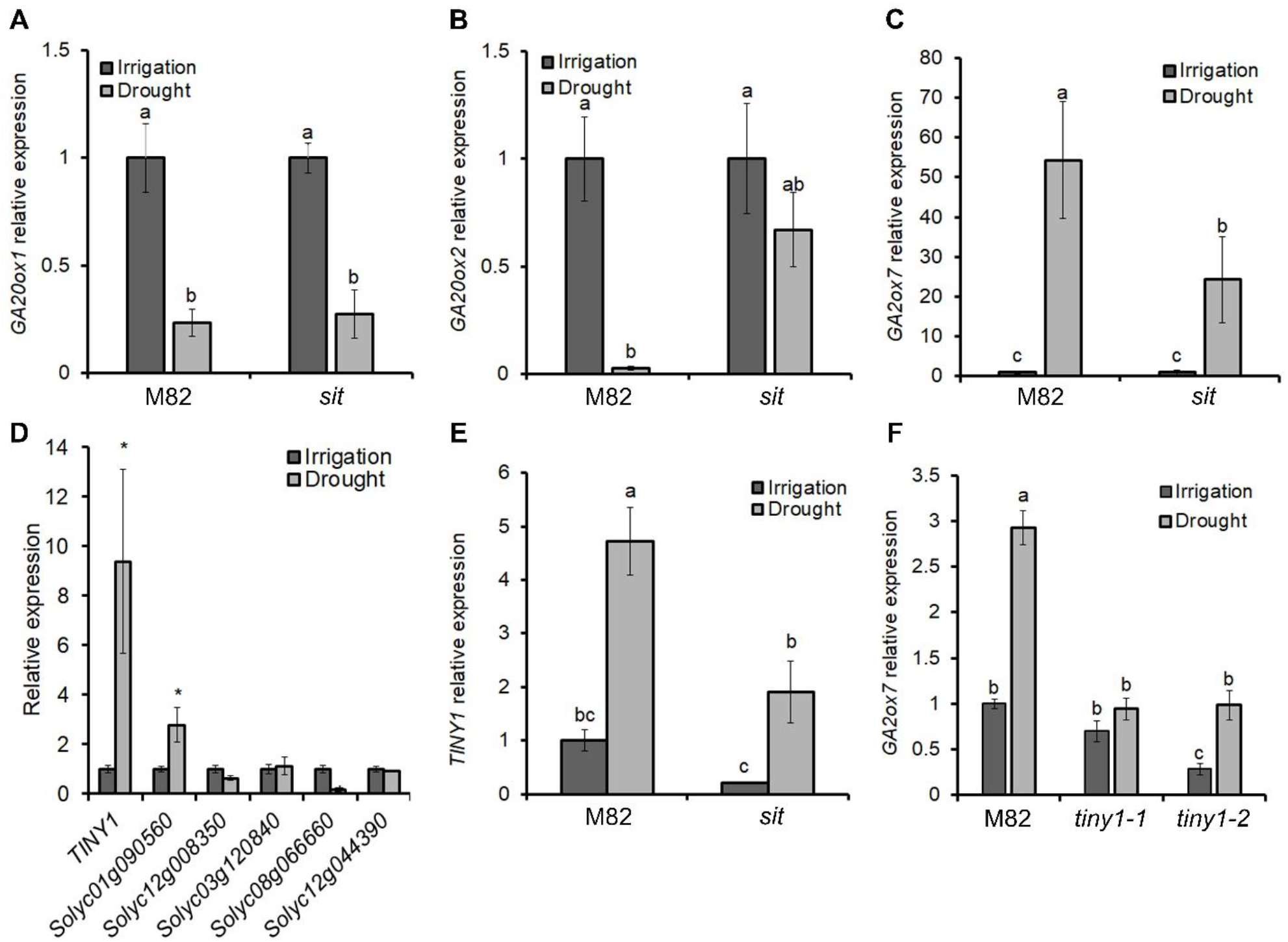
Drought inhibition of GA accumulation is partially mediated by the ABA-induced *TINY1*. **(A), (B)** and **(C)** Relative expression of *GA20ox1* **(A)**, *GA20ox2* **(B)** and *GA2ox7* **(C)** under irrigation and drought conditions (Soil VWC 15%) in leaves of M82 and the ABA-deficient mutant *sit*. Values are means of 4 biological replicates ±SE. **(D)** Relative expression of the six putative *TINY* genes in leaves of M82 plants grown with irrigation or exposed to drought condition (Soil VWC 15%). **(E)** Relative expression of *TINY1* in leaves of M82 and the ABA-deficient mutant *sit* grown with irrigation or exposed to drought condition (Soil VWC 15%). Values are means of 4 biological replicates ±SE. **(F)** Relative expression of *GA2ox7* under irrigation and drought conditions in leaves of M82 and *tiny1-1* and *tiny1-2*. Values are means of 4 biological replicates ±SE. Small letters **(A, B, C, E** and **F)** or stars **(D)** represent significant differences between respective treatments by Tukey-Kramer HSD test (*P < 0*.*05)* or Student’s *t* test (*P* < *0*.*05*), respectively.

A previous study in tomato suggested a role for a DREB transcription factor from the sub-family TINY in GA metabolism (Li et al., 2012). *In silico* analysis of *GA20ox1, GA20ox2* and *GA2ox7* promoters suggested the presence of several putative DREB-binding elements (Sakuma et al., 2002) in *GA2ox7*, but not in *GA20ox1* and *GA20ox2* promoters (Supplemental Fig. 6). Tomato has 6 putative DREB-TINY genes. Only *TINY1* (Solyc06g066540) and Solyc01g090560 were upregulated by drought (Fig. 2D) and the effect on *TINY1* was much stronger. The upregulation of *TINY1* by water-deficit conditions was partially inhibited in *sit* (Figure 2E) and the application of ABA increased its expression (Supplemental Figure 5B), suggesting that drought-induced *TINY1* expression, is partially ABA-dependent, similar to *GA2ox7*. To examine if *TINY1* affects the expression of *GA2ox7*, we generated CRISPR-Cas9 derived *tiny1* mutants (Supplemental Fig. 7). The homozygous mutant lines (two alleles, *tiny1-1* and *tiny1-2*) showed a WT phenotype. The loss of *TINY1* had no effect on *GA20ox1* and *GA20ox2* downregulation by drought (Supplemental Fig. 5C and D). However, drought-induced *GA2ox7* expression was strongly inhibited in *tiny1-1* and *tiny1-2* (Fig. 2F). Together, the results imply that drought-induced *GA2ox7* expression is regulated (directly or indirectly) by *TINY1*.

### GA deactivation in guard cells promotes stomatal closure under water-deficit conditions

Since water deficiency upregulated *GA2ox7* expression in guard cells, we examined the significance of GA2ox7 activity to stomatal closure. *ga2ox7* mutant was recently characterized in tomato (Schrager-Lavelle et al., 2019). The mutant has elongated epicotyl but normal leaves with WT stomatal density (Supplemental Fig.8, Fig. 3A). Under normal irrigation, the loss of *GA2ox7* had no effect on the basal stomatal aperture, transpiration rate and stomatal conductance (Supplemental Fig. 9). However, when *ga2ox7* plants were exposed to water-deficit conditions, they exhibited reduced ‘drought avoidance’; they wilted before the WT and exhibited a faster decrease in leaf relative water content (RWC, Fig. 3A and B). We then examined if the rapid water loss was caused by a higher transpiration rate under water-deficit conditions. We analyzed whole-plant transpiration in WT and *ga2ox7* mutant plants, grown in a greenhouse using an array of lysimeters (Illouz-Eliaz et al., 2020). Plants were grown with irrigation, and then water supply was terminated. Transpiration rate was not affected in the first three days into the drought treatment (Fig. 3C). However, on the fourth day, the transpiration rate in WT plants sharply decreased, but that of *ga2ox7* did not change. Only on the fifth day, *ga2ox7* plants reduced their transpiration. Whole-plant daily transpiration was also decreased in WT before *ga2ox7* (Supplemental Fig. 10A). We further analyzed the transpiration rate during gradual soil dehydration by thermal imaging. In well-watered soil (60% soil VWC) leaf surface temperature was similar in WT and *ga2ox7*, indicating a similar transpiration rate (Fig. 3D). However, when the soil was dehydrated to 35% VWC, the temperature of WT leaves increased, but that of *ga2ox7* did not change, indicating that WT but not the mutant closed their stomata. At 15% soil VWC, WT and *ga2ox7* leaves exhibited higher and similar temperature, indicating that both closed their stomata. Microscopic analysis showed that under mild water deficiency (35% soil VWC) 70% of WT stomata were closed while only 49% in *ga2ox7* (Fig. 3E). Finally, we analyzed stomatal conductance in WT and *ga2ox7* plants at different soil VWC. In irrigated plants, stomatal conductance was similar in WT and the mutant (Fig. 3F, Supplemental Fig. 10B). However, when soil VWC was reduced to 40%, stomatal conductance in WT was significantly lower than in *ga2ox7*. Under severe drought (15% soil VWC), stomatal conductance was very low and similar in WT and *ga2ox7*. These results show that *ga2ox7* stomata are hyposensitive to soil dehydration. Analysis of *GA2ox7* expression in guard cells during soil dehydration showed upregulation already at the early stages of soil dehydration (30% soil VWC, Fig. 3G). Taken together, the results suggest that GA deactivation at the early stages of soil dehydration contributes to stomatal closure.

**Figure 3.**
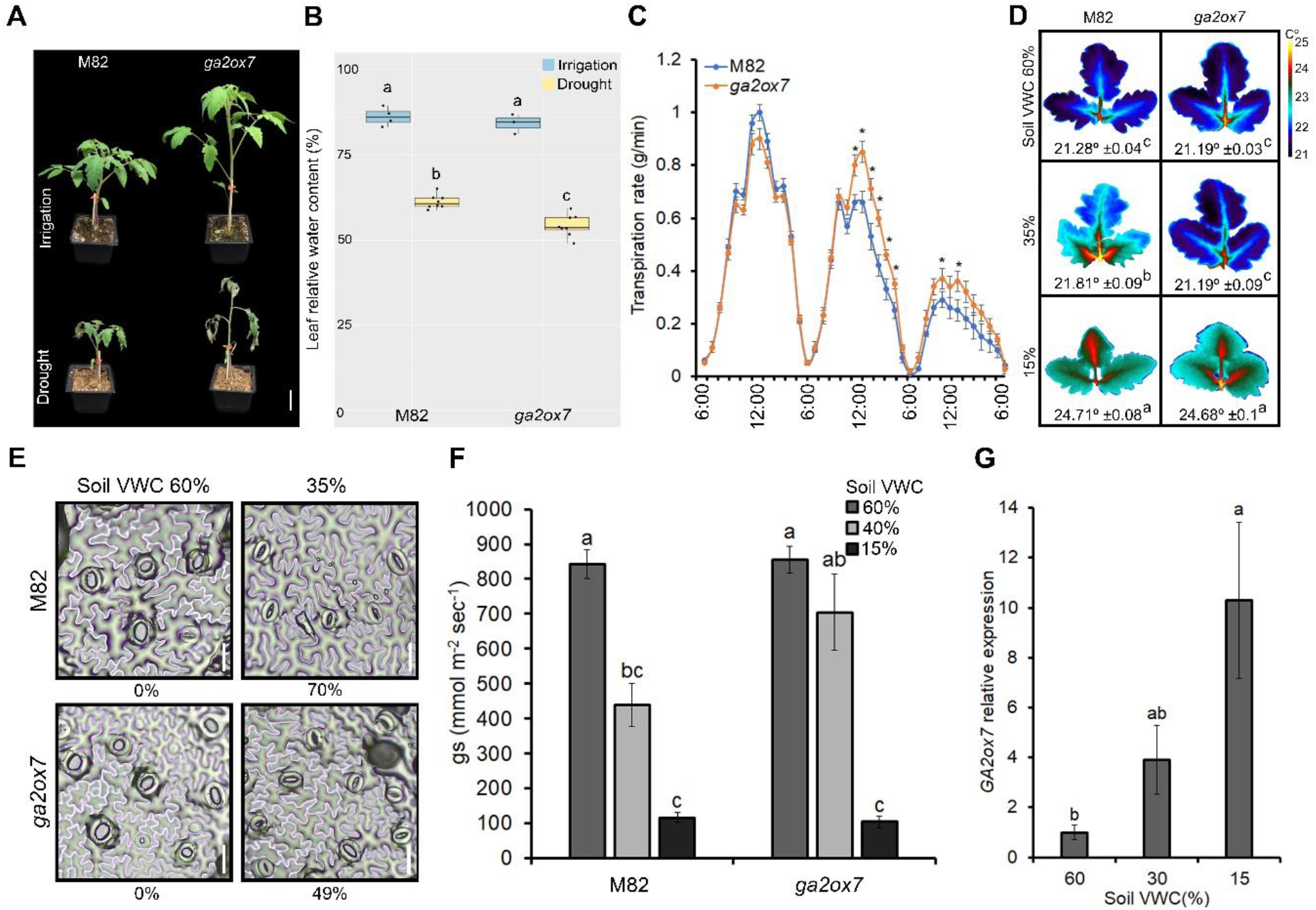
*ga2ox7* stomata are hyposensitive to soil dehydration. **(A)** Representative plants grown under irrigation or 7 days without irrigation. Scale bar = 5 cm. **(B)** Leaf relative water content (RWC) in M82 and *ga2ox7* grown under irrigation or without irrigation for 7 days. Values are means of 4 (for irrigation) or 8 (for drought) biological replicates ±SE. **(C)** M82 and *ga2ox7* whole-plant transpiration rate over the course of 24h (6:00 AM to 6:00 AM) 3 to 5 days after the termination of irrigation. Values are means of 6 (for M82) or 10 (for *ga2ox7*) plants ±SE. M82 and *ga2ox7* plants were placed on lysimeters and pot (pot + soil + plant) weight was measured every 3 min. **(D)** Thermal imaging of representative leaves (leaf # 4 below the apex) of M82 and *ga2ox7* plants exposed to different soil VWC. Images were digitally extracted for comparison. Numbers below leaves are the average leaf-surface temperature, and the values are means of 5 biological replicates (plants), each measured 6 times ± SE. **(E)** Representative abaxial epidermal imprints taken from irrigated or drought (mild drought, soil VWC 35%) treated M82 and *ga2ox7*. Numbers below images represent the percentage of closed stomata in each treatment. Values are percentages of 4 biological replicates each with approximately 100 measurements (stomata). Scale bar = 30 µm. **(F)** Stomatal conductance (g_s_) of M82 and *ga2ox7* under different soil VWC. Values are means of 4 biological replicates ±SE. **(G)** *GA2ox7* relative expression in M82 isolated guard cells under different soil VWC. Stars or small letters represent significant differences between respective lines (C) by Student’s *t* test (*P* < *0*.*05*) or lines and treatments **(B, D, F** and **G)** Tukey-Kramer HSD test (*P* < 0.05).

### Mutations in *GA20ox1* and *GA20ox2* promoted ‘drought avoidance’ by suppressing leaf expansion

To test if the reduced expression of *GA20ox1* and *GA20ox2* also affects stomatal closure and water status under water-deficit conditions, we generated CRISPR/Cas9-derived *ga20ox1* and *ga20ox2* loss-of-function mutants (Supplemental Fig. 11). Both mutants exhibited shorter stems, smaller leaves but WT basal stomatal aperture in well-watered plants (Fig. 4A; Supplemental Fig. 12). We first examined ‘drought avoidance’ in these two mutants. To this aim, we exposed WT, *ga20ox1* and *ga20ox2* plants to water-deficit conditions and analyzed the rate of water loss. Both mutants maintained higher leaf RWC compared to WT (Fig. 4B), indicating a reduced rate of water loss under drought. We then analyzed the transpiration rate using thermal imaging. Both mutations had no effect on transpiration rate under irrigation or water-deficit conditions (Fig, 4C). In addition, stomatal conductance was similar between WT and the mutants during soil dehydration (Fig. 4D). Together, the results suggest that the reduced rate of water loss found in *ga20ox1* and *ga20ox2* plants under water-deficit conditions was caused by the smaller leaf area, and not by faster stomatal closure. We also generated the double mutant *ga20ox1/ga20ox2* by crosses. The double mutant exhibited an additive effect on growth; plants were dwarf and their leaves were very small (Supplemental Fig. 13). Microscopic analysis of *ga20ox1/ga20ox2* abaxial leaf epidermis showed reduced stomatal aperture (Fig. 4E), similar to other strong GA mutants, such as overexpression of stable DELLA (Nir et al., 2017). In line with this observation, stomatal conductance in the double mutant was lower than WT in well-watered plants (Fig. 4F). These results suggest high redundancy between *GA20ox*1 and *GA20ox2* in the promotion of stomatal closure, but not in the regulation of leaf growth. *GA20ox1* and *GA20ox2* expression was much higher in whole-leaf tissue than in guard cells (Fig. 4G). This may explain why they have a significant role in leaf growth but not in stomatal closure.

**Figure 4.**
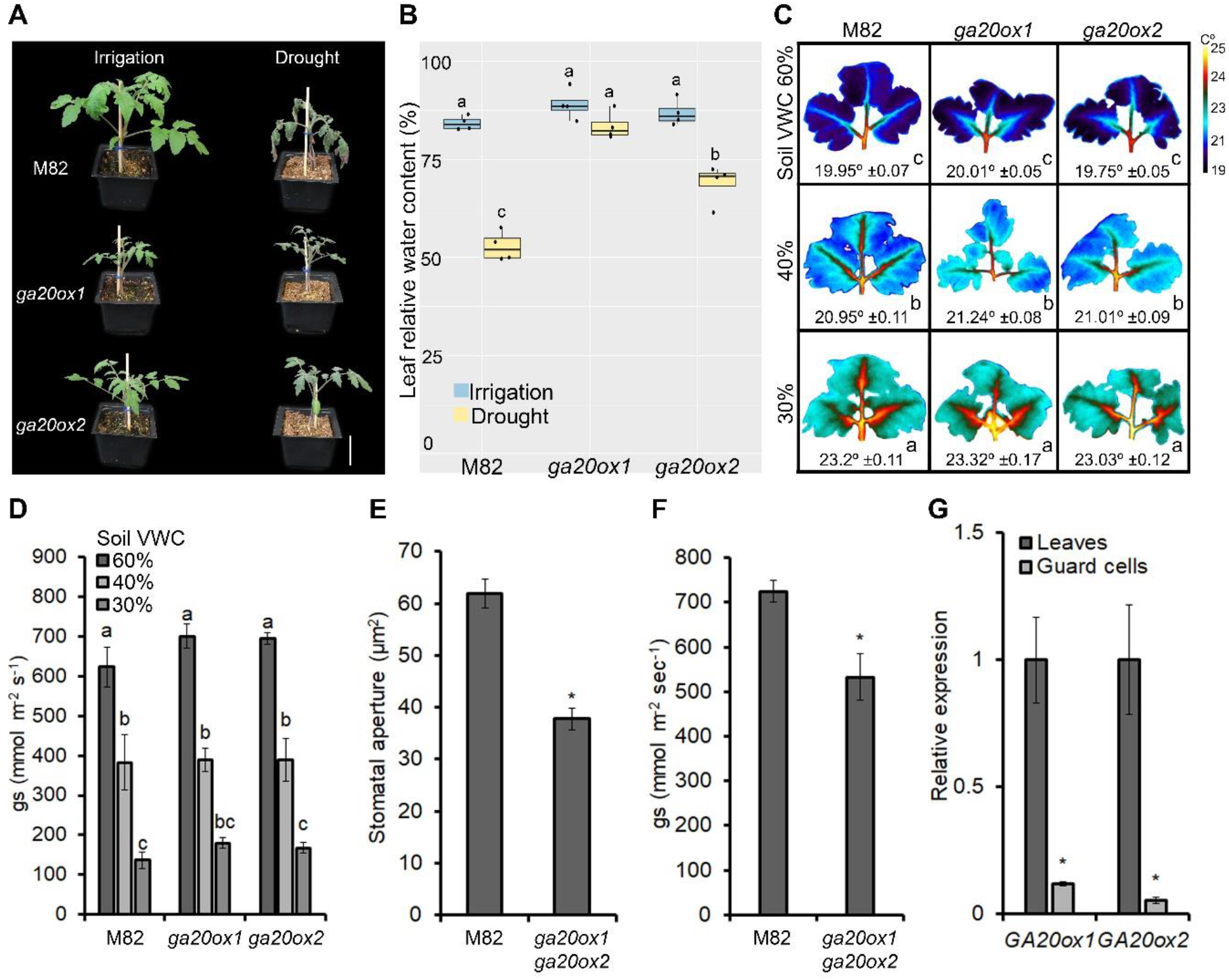
Loss of *GA20ox1* and *GA20ox2* reduced canopy growth and inhibited water loss under water-deficit conditions. **(A)** Representative plants grown under irrigation or 7 days of drought. Scale bar = 5 cm. **(B)** Leaf RWC in M82, *ga20ox1* and *ga20ox2* grown under irrigation or 7 days without irrigation. Values are means of 4 biological replicates ±SE. **(C)** Thermal imaging of representative leaves of M82, *ga20ox1* and *ga20ox2* plants exposed to different soil VWC. Images were digitally extracted for comparison. Numbers below plants are the average leaf-surface temperature, and the values are means of 5 biological replicates (plants), each measured 6 times ± SE. **(D)** Stomatal conductance (g_s_) in M82, *ga20ox1* and *ga20ox2* under different soil VWC. Values are means of 4 biological replicates measured 4 to 8 times ±SE. **(E)** Stomatal aperture of M82 and the double mutant *ga20ox1/ga20ox2* measured on imprints of abaxial epidermis taken from leaf # 3 below the apex. Values are means of 4 biological replicates each with approximately 100 measurements (stomata) ±SE. **(F)** Stomatal conductance (g_s_) of M82 and *ga20ox1*/*ga20ox2*. Values are means of 4 biological replicates ±SE. **(G)** Relative expression of *GA20ox1* and *GA20ox2* in whole-leaf tissue compared to isolated guard cells. Values are means of 4 biological replicates ±SE. Small letters **(B, C** and **D)** or stars **(E, F** and **G)** represent significant differences between respective treatments by Tukey-Kramer HSD test (*P < 0*.*05)* or Student’s *t* test (*P* < *0*.*05*), respectively.

We further generated a double mutant *ga20ox1/ga2ox7* by crosses. The homozygous double mutant plants exhibited elongated epicotyl, similar to *ga2ox7*, and smaller leaves, similar to *ga20ox1* (Supplemental Fig. 14, Fig. 5A and B). While *ga2ox7* single mutant exhibited rapid wilting and low RWC under water-deficit conditions, *ga20ox1* plants maintained higher leaf RWC and showed slower water loss. In the double mutant, the loss of *GA20ox1*, strongly suppressed the effect of *ga2ox7*; these plants maintained higher leaf RWC, slightly lower than the single mutant *ga20ox1* (Fig. 5B and C). These results demonstrate the importance of leaf area to ‘drought avoidance’.

**Figure 5.**
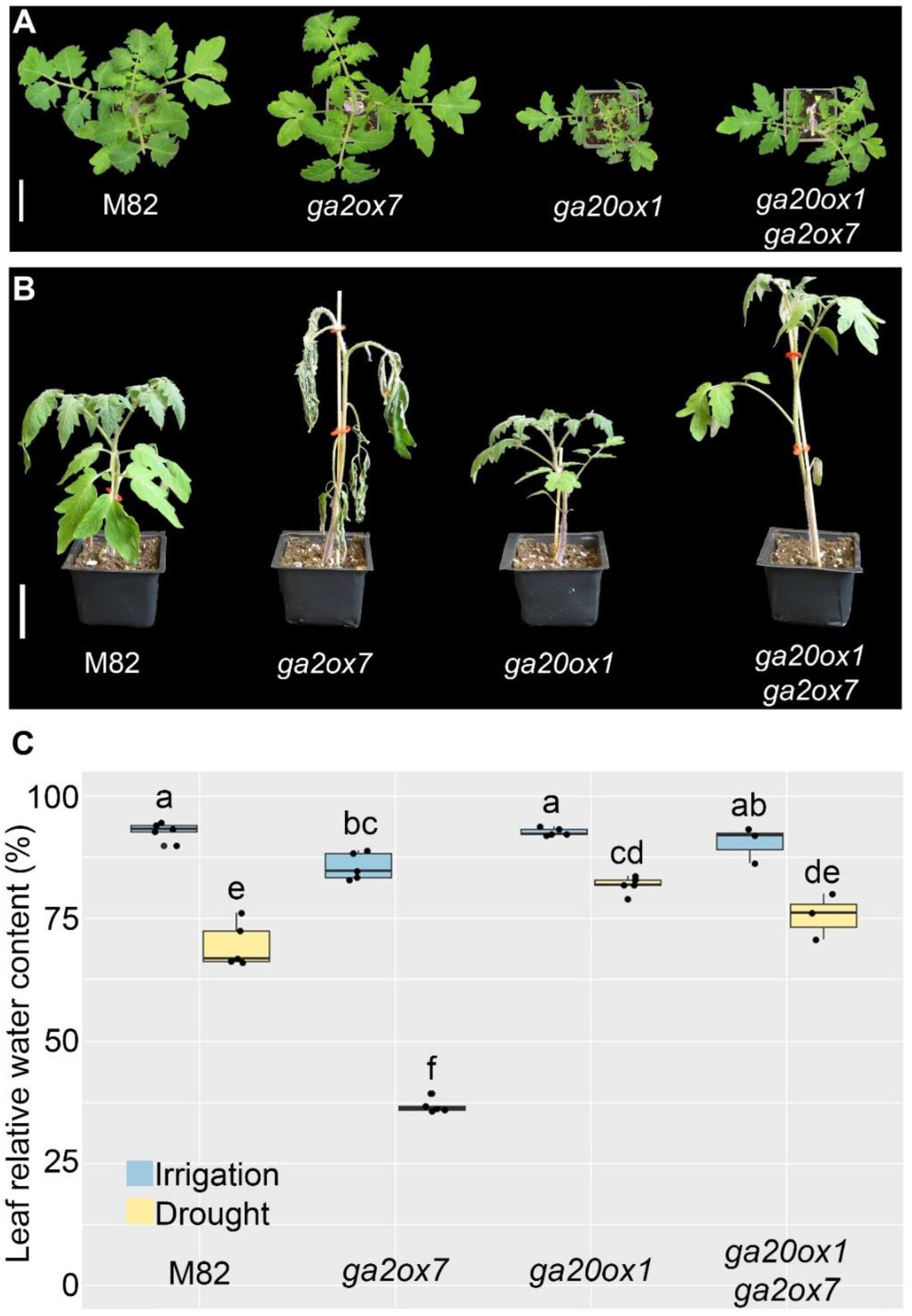
Reduced GA levels contribute to ‘drought avoidance’ by suppressing leaf growth and promoting stomatal closure. **(A)** and **(B)** Representative plants grown under irrigation **(A**) or 10 days without irrigation **(B)**. Scale bar = 5 cm. **(C)** Leaf RWC in M82, *ga2ox7, ga20ox1* and the double mutant *ga20ox1/ga2ox7* under irrigation or 10 days without irrigation. Values are means of 3-5 biological replicates (four terminal leaflets taken from leaf # 3 below the apex) ±SE. Small letters **(C)** represent significant differences between respective treatments by Tukey-Kramer HSD test (*P < 0*.*05)*.

## DISCUSSION

‘Drought avoidance’ is a major plant adaptation strategy to survive transient water-deficit conditions (Skirycz and Inze, 2010; Kooyers, 2015). To avoid drought stress, plants reduce their transpiration and can use the available water in the soil more slowly and for a longer period before the arrival of the next rain. Two major mechanisms have been evolved to reduce water loss under drought; fast stomatal closure and long-term growth inhibition. Here we demonstrate the role of the growth-promoting hormone GA in the regulation of these fast and long-term drought responses.

Previous studies showed that GA treatments promote stomatal opening, whereas reduced GA activity by overexpressing stable DELLA, accelerates stomatal closure (Santakumari and Fletcher, 1987; Göring et al., 1990; Nir et al., 2017). The results of this study suggest that water deficiency reduces the levels of the bioactive GA_1_ and GA_3_ in tomato guard cells and this accelerates stomatal closure. The relatively high levels of GA_3_ in stomata of well-watered plants were unexpected since most studies found only traces amount of this bioactive GA in plants (Hedden, 2020). However, a recent study showed the accumulation of GA_3_ in tomato fruits (Li et al., 2020). The reduced levels of bioactive GAs in guard cells under water deficiency is probably a result of the strong upregulation of the GA-deactivating gene *GA2ox7. GA2ox7* belongs to class III *GA2oxs*. Enzymes of this class catalyze the deactivation of C-20 GA precursors, affecting the metabolic flow towards the production of C-19 bioactive GAs (Hedden 2020). Water deficiency reduced in guard cells not only the level of the bioactive GAs but also the level of their C-20 precursor GA_12_, the direct substrate of GA2ox7 (Schrager-Lavelle et al., 2019). *GA2ox7* was upregulated in guard cells at the early stages of soil dehydration and its loss-of-function inhibited stomatal closure in response to mild soil dehydration. Under severe drought, however, stomata of *ga2ox7* were closed, similar to those of WT. Previously we showed that reduced GA activity promotes ABA responses in guard cells (Nir et al., 2017; Shohat et al., 2020). Promoting ABA responses can have a significant effect on stomatal closure at the early stages of soil dehydration when ABA levels are still limited, but in dry soil, when ABA levels are saturated, the effect is probably neglected. Thus, we propose that at the early stages of soil dehydration, *GA2ox7* is upregulated in guard cells, leading to a reduction in bioactive-GA levels that in turn, increases ABA activity and accelerates stomatal closure (Fig. 6).

**Figure 6.**
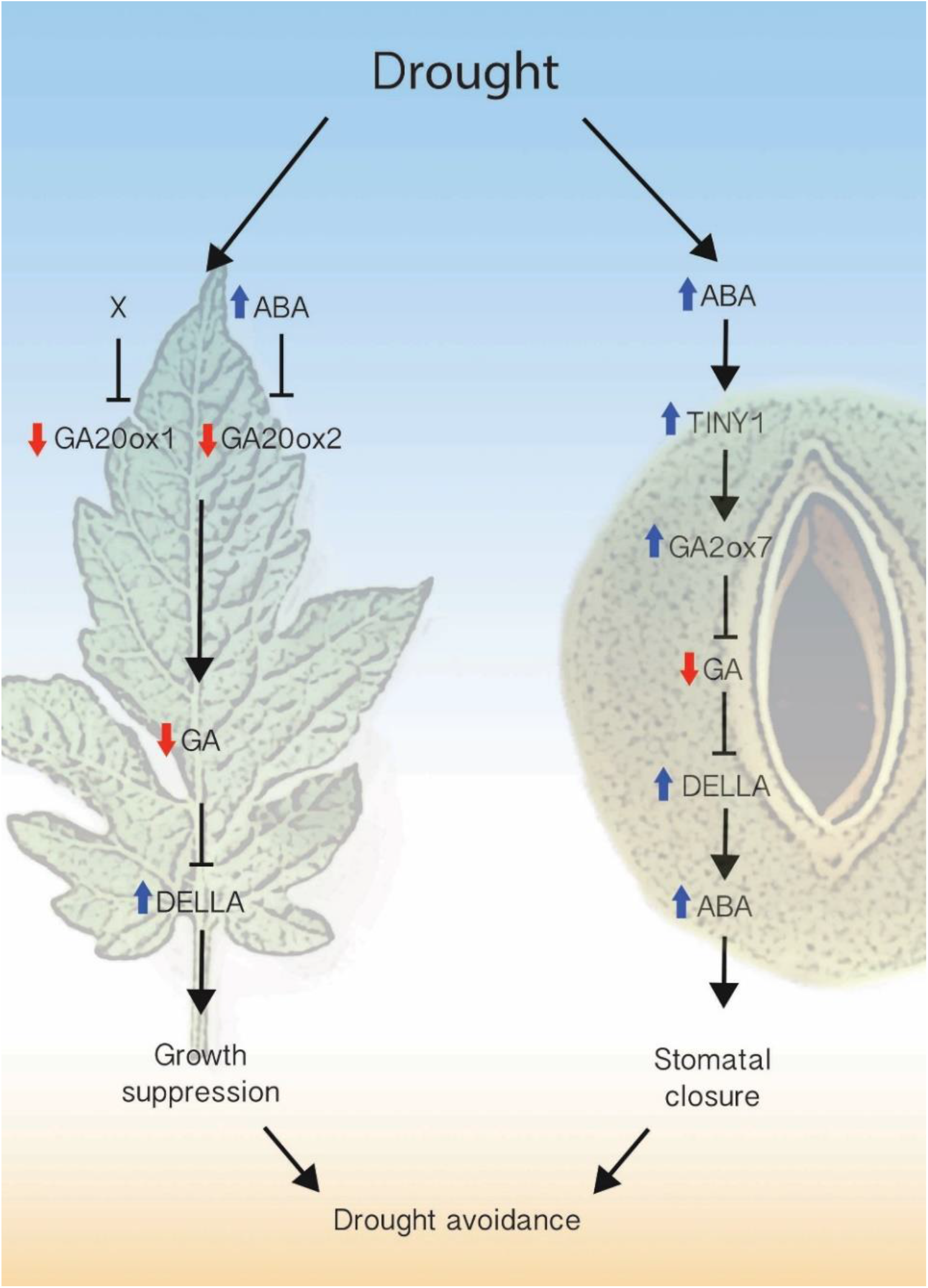
Suggested model for the role of GA in ‘drought avoidance’ in tomato plants. Water-deficit conditions, via ABA-dependent and independent pathways downregulates the expression of the GA biosynthesis genes *GA20ox1* and *GA20ox2* and upregulates the GA deactivating gene *GA2ox7* in leaf tissue and guard cells. The upregulation of *GA2ox7* by drought and ABA is mediated by the transcription factor DREB-TINY1. These molecular changes lead to reduced levels of bioactive GAs and the accumulation of DELLA. In turn, DELLA inhibits leaf growth and promotes ABA-induced stomatal closure at the early stages of soil dehydration. In leaf tissue the levels of *GA20ox1* and *GA20ox2* expression has the dominant role in the regulation of growth, whereas in guard cells *GA2ox7* has a major role in stomatal closure. The inhibition of leaf growth and the earlier stomatal closure reduce transpiration and promote ‘drought avoidance’.

It is not clear yet what is the contribution of *GA20ox1* and *GA20ox2* activity to these changes in guard-cell GA levels. The expression of these two genes in guard cells was suppressed under drought, but their loss-of-function had no effect on basal stomatal aperture or stomatal response to water deficiency. It had a strong effect, however, on leaf growth. Only in the double knockout mutant *ga20ox1/ga20ox2* the basal stomatal aperture and conductance were reduced, similar to overexpression of stable DELLA (Nir et al., 2017). Since these two genes exhibited low expression in guard cells (compared to whole-leaf tissue), their contribution to GA accumulation in these cells may be limited. It is possible that GA levels/activity in guard cells has to be below a certain threshold to affect stomatal closure.

*GA20ox1* and *GA20ox2* expression was suppressed by water deficiency also in leaf tissues. Although the single mutants exhibited WT stomatal closure, both showed reduced water loss during water-deficit treatment and improved ‘drought avoidance’. This effect can be attributed to the reduced leaf size. Water-deficit conditions suppress growth to reduce transpiration area and relocate resources for adaptation (Eziz et al., 2017). It was shown previously that growth inhibition by salt stress is mediated by reduced GA accumulation (Achard et al., 2006). Our results in tomato, suggest that growth inhibition by drought is also mediated, at least partially, by a reduction in bioactive GA levels. The suppression of leaf growth seems to contribute significantly to the reduction in whole-plant transpiration and ‘drought avoidance’. In the *ga2ox7/ga20ox1* double mutant, the effect of reduced leaf size, caused by the loss of *GA20ox1*, suppressed the fast water loss caused by *ga2ox7*. It is interesting to note that the double mutant *ga2ox7/ga20ox1* exhibits smaller leaves, similar to *ga20ox1*, and elongated epicotyl, similar to *ga2ox7*. This suggest that the high GA levels in the stem cannot compensate for the low GA levels in the leaves, and implies that GAs are not translocated from stems to leaves in tomato.

The expression of *GA20ox* genes is regulated by the GA-feedback loop via the transcriptional complex DELLA-INDETERMINATE DOMAIN (IDD, Fukazawa et al., 2017) to maintain GA homeostasis. A major question is how environmental conditions overcome this mechanism of homeostasis. In tomato leaves, water deficiency inhibited the expression of *GA20ox1* and *GA20ox2*, but induced the expression of *GA20ox3*. These three genes (*GA20ox1/2/3*) were upregulated in transgenic plants overexpressing stable DELLA protein (*proΔ17*), suggesting that all of them are regulated by the feedback response via DELLA to maintain GA homeostasis. Water deficiency, however, downregulated *GA20ox1* and *GA20ox2* (but not *GA20ox3*) in the presence of stable DELLA (*proΔ17*) and inhibited the accumulation of bioactive GAs. These suggest that water deficiency out-competes the effect of DELLA on the transcriptional upregulation of the two genes, and keeps them low despite DELLA accumulation and therefore, suppresses the mechanism of homeostasis.

Plant responses to drought and ABA are mediated by the ETHYLEN RESPONSE FACTOR (ERF)/AP2 transcription factors DREB (Sakuma et al., 2002). DREBs regulate downstream responses, including stomatal closure, growth suppression and induction of stress-related genes (Lata and Prasad, 2011). DREB proteins are divided into six subfamilies (A1 to A6), and DREB-TINYs belong to subfamily A4, which contains 16 genes in Arabidopsis (Nakano et al., 2006). An Arabidopsis TINY protein suppresses growth by inhibition of brassinosteroid activity and promotes ABA-induced stomatal closure (Xie et al., 2019). In tomato, overexpression of DREB-*TINY1* suppresses *GA20ox1* and *GA20ox2* expression, reduces GA levels, inhibits growth, and promotes tolerance to drought (Li et al., 2012). The tomato *TINY1* was also induced by high temperatures and its downregulation resulted in susceptibility to heat stress (Mao et al., 2020). Our results show that the loss of *TINY1* had no effect on the suppression of *GA20ox1* and *GA20ox2* by drought, suggesting that TINY1 is not the drought-induced regulator of these *GA20oxs*. On the other hand, *GA2ox7* induction by drought was inhibited in *tiny1*, suggesting that *TINY1* mediates the effect of drought on *GA2ox7* expression. Whether *TINY1* is a direct regulator of *GA2ox7*, is not yet clear.

To conclude, we suggest that water-deficit conditions, via ABA-dependent and independent pathways, downregulate the expression of the GA biosynthesis genes *GA20ox1* and *GA20ox2* and upregulate the GA deactivating gene *GA2ox7* in leaf tissues and guard cells, leading to reduced levels of bioactive GAs. The lower GA levels in leaves suppress their growth and in guard cells promote ABA-induced stomatal closure at the early stages of soil dehydration (Fig. 6). The suppression of leaf growth is regulated mainly by the inhibition of GA biosynthesis (downregulation of *GA20ox*), whereas the accelerated stomatal closure, by GA deactivation (upregulation of *GA2ox*). These short-and long-term responses reduce transpiration and promote ‘drought avoidance’ and adaptation to water-deficit conditions.

## METHODS

### Plant materials, growth conditions and hormone treatments

Tomato (*Solanum lycopersicum*) plants in M82 background (*sp/sp*) were used throughout this study. The *ga2ox7, sit*, transgenic line 35S:*proΔ17* and the CRISPER-derived *ga20ox1, ga20ox2* and *tiny1* were backcrossed to or generated in M82 background. Plants were grown in a growth room set to a photoperiod of 12/12-h night/day, light intensity of 150 μmol m^-2^ s^-1^ and 25°C and irrigated to saturation. In other experiments, plants were grown in a greenhouse under natural day-length conditions, light intensity of 700 to 1000 µmol m^-2^ s^-1^ and 18-30°C. The seeds were harvested from ripe fruits and treated with 1% sodium hypochlorite followed by 1% Na_3_PO_4_ 12H_2_O, and incubated with 10% Sucrose over-night at 37°C. Seeds were stored dry at room temperature. (±)-ABA dissolved in DMSO (Sigma-Aldrich, St. Louis, USA) was applied to plants by spraying.

### CRISPR/Cas9 mutagenesis: cloning, plant transformation and selection of mutant alleles

Two single-guide RNAs (sgRNAs, Table S1) were designed to target *GA20ox1, GA20ox2* and *TINY1* genes, using the CRISPR-P tool (http://cbi.hzau.edu.cn/crispr). Vectors were assembled using the Golden Gate cloning system, as described by Weber et al. (2011). Final binary vectors, pAGM4723, were introduced into *A. tumefaciens* strain GV3101 by electroporation. The constructs were transformed into M82 cotyledons using transformation and regeneration methods described by McCormick (1991). Kanamycin-resistant T0 plants were grown and independent transgenic lines were selected and self-pollinated to generate homozygous transgenic lines. The genomic DNA of each plant was extracted, and genotyped by PCR for the presence of the Cas9 construct. The CRISPR/Cas9-positive lines were further genotyped for mutations using a forward primer to the upstream sequence of the sgRNA1 target and a reverse primer to the downstream of the sgRNA2 target sequence. The target genes in all mutant lines were sequenced. Several homozygous and heterozygous lines were identified and independent mutant lines for each gene were selected for further analysis. The Cas9 construct was segregated out by crosses to M82.

### Isolation of guard cells

Guard cells from tomato leaves (leaves no. 3 and 4 below the apex) were isolated according to Shohat et al. (2020). Briefly, four fully expanded leaves without the central veins were ground twice in a blender containing 100 ml cold distilled water, for 30 sec each time. The blended mixture was poured onto a 100 µm nylon mesh (Sefar) and the remaining epidermal peels were rinsed thoroughly with 0.5L of cold deionized water. The peels were then transferred into 2 ml Eppendorf tubes and frozen in liquid nitrogen. The samples were stained with 0.03% Neutral red and cell vitality was examined under light microscope.

### RNA extraction and cDNA synthesis

Total was RNA extracted by RNeasy Plant Mini Kit (Qiagen). For synthesis of cDNA, SuperScript II reverse transcriptase (18064014; Invitrogen, Waltham, MA, USA) and 3 mg of total RNA were used, according to the manufacturer’s instructions.

### RT-qPCR analysis

RT-qPCR analysis was performed using an Absolute Blue qPCR SYBR Green ROX Mix (AB-4162/B) kit (Thermo Fisher Scientific, Waltham, MA 15 USA). Reactions were performed using a Rotor-Gene 6000 cycler (Corbett Research, Sydney, Australia). A standard curve was obtained using dilutions of the cDNA sample. The expression was quantified using Corbett Research Rotor-Gene software. Three independent technical repeats were performed for each sample. Relative expression was calculated by dividing the expression level of the examined gene by that of *SlACTIN*. Gene to *ACTIN* ratio was then averaged. All primer sequences are presented in Supplemental Table 2. The values for control (Mock and irrigation) and/or M82 wild type treatments was set to 1.

### GA analysis in leaves

The sample preparation and analysis of gibberellins (GAs) was performed according to the method described in Urbanová et al. (2013) with some modifications. Briefly, tissue samples of about 10 mg DW were ground to a fine consistency using 3-mm zirconium oxide beads (Retsch GmbH & Co. KG, Haan, Germany) and a MM 301 vibration mill at a frequency of 30 Hz for 3 min (Retsch GmbH & Co. KG, Haan, Germany) with 1 mL of ice-cold 80 % acetonitrile containing 5 % formic acid as extraction solution. The samples were then extracted overnight at 4 °C using a benchtop laboratory rotator Stuart SB3 (Bibby Scientific Ltd., Staffordshire, UK) after adding 17 internal gibberellins standards ([^2^H_2_]GA_1_, [^2^H_2_]GA_3_, [^2^H_2_]GA_4_, [^2^H_2_]GA_5_, [^2^H_2_]GA_6_, [^2^H_2_]GA_7_, [^2^H_2_]GA_8_, [^2^H_2_]GA_9_, [^2^H_2_]GA_15_, [^2^H_2_]GA_19_, [^2^H_2_]GA_20_, [^2^H_2_]GA_24_, [^2^H_2_]GA_29_, [^2^H_2_]GA_34_, [^2^H_2_]GA_44_, [^2^H_2_]GA_51_ and [^2^H_2_]GA_53_; purchased from OlChemIm, Czech Republic. The homogenates were centrifuged at 36 670 *g* and 4 °C for 10 min, corresponding supernatants further purified using reversed-phase and mixed-mode SPE cartridges (Waters, Milford, MA, USA) and analyzed by ultra-high-performance liquid chromatography-tandem mass spectrometry (UHPLC-MS/MS; Micromass, Manchester, UK). GAs were detected using multiple-reaction monitoring mode of the transition of the ion [M–H]^-^ to the appropriate product ion. Masslynx 4.1 software (Waters, Milford, MA, USA) was used to analyze the data and the standard isotope dilution method (Rittenberg and Foster, 1940) was used to quantify the GAs levels.

### GA analysis in guard cells

Hormone extraction was performed as described previously described (Breitel et al., 2016; Kojima et al., 2009) with some modifications. Briefly, isolated guard cells samples were frozen in liquid nitrogen and grounded into powder using motor and pestle. Gibberellins were extracted from 200 mg of the ground sample in ice-cold methanol/water/formic acid (15/4/1 v/v/v) added with internal standards. Similar concentrations of isotope labeled gibberellin internal standards were added into samples and calibration standards. The samples were then purified using Oasis MCX SPE cartridges (Waters) according to manufacturer’s protocol and injected on Acquity UPLC BEH C18 column (1.7 µm, 2.1×100 mm, Waters; mobile phases: gradients of 0.1% acetic acid in water or acetonitrile), connected to Acquity UPLC H class system (Waters) coupled with UPLC-ESI-MS/MS triple quadrupole mass spectrometer for identification followed by quantification of hormones. The hormones were measured in negative mode, with two MRM transitions for each compound. External calibration curves were constructed with a series of diluted gibberellin standards and deuterium labeled internal standards (Table S3) and used for absolute quantification (Balcke et al., 2012). Hormone concentrations were derived by comparing ratios of MRM peak areas of analyte with its corresponding internal standard using Target Lynx (v4.1; Waters) software.

### Thermal imaging

Thermal images were obtained using an A655SC, FOV 15 (FLIR Systems, Wilsonville, USA). The camera was mounted vertically above the plants. Mean temperature of leaflets from leaves no. 3 and 4 below the apex were calculated using the customized region of interest (ROI) tool, according to the manufacturer’s instructions.

### Stomatal aperture measurements

Stomatal aperture was determined using the rapid imprinting technique described by Geisler *et al*. (2000). Light-bodied vinylpolysiloxane dental resin (eliteHD+, Zhermack Clinical, Badia Polesine, Italy) was attached to the abaxial side of the leaflet and then removed as soon as it dried (minutes). The resin epidermal imprints were covered with transparent nail polish, which was removed once it dried and served as a mirror image of the resin imprint. The nail-polish imprints were put on glass cover-slips and photographed under a model ICC50 W bright-field inverted microscope (Leica Microsystem, Wetzlar, Germany). Stomatal images were later analyzed to determine aperture size using the ImageJ software fit-ellipse tool (http://rsb.info.nih.gov/ij/). A microscopic ruler (Olympus) was used for size calibration.

### Transpiration rate and daily transpiration measurements

Whole-plant transpiration rate was determined using an array of lysimeters placed in the greenhouse (Plant array 3.0 system; Plant-Ditech) in the I CORE Center for Functional Phenotyping (http://departments.agri.huji.ac.il/plantscience/icore.phpon), as described in detail by Halperin et al. (2017). Briefly, plants were grown in 4-liter pots under semi-controlled temperature conditions (20/32 °C night/day), natural day-length, and light intensity of approximately 1000 µmol m^−2^ s^−1^. Each pot was placed on a temperature-compensated load cell with digital output (Vishay Tedea-Huntleigh) and sealed to prevent evaporation from the surface of the growth medium. The weight output of the load cells was monitored every 3 min. The data were analyzed using SPAS analytics (Plant-Ditech) software to obtain the following whole-plant physiological traits: daily transpiration (weight loss between predawn and sunset) and transpiration rate (weight loss between two 3-min time points) were calculated from the weight difference between the two data points.

### Stomatal conductance (g_s_) measurements

Stomatal conductance was determined using a SC-1 Leaf Porometer (Decagon Devices, Pullman, WA, USA) or a LI-6800 portable gas exchange system (LI-COR Biosciences). Measurements were performed at 9:00 AM under a constant CO_2_ concentration of 400 ppm and constant photosynthetic photon flux density (PPFD) of 1200 µmol m^−2^ s^−1^.

### Measurements of leaf relative water content (RWC)

Leaf RWC of irrigated and drought-treated plants were measured as follows: Fresh weight (FW) was measured immediately after leaf detachment and then leaves were soaked for 24h in 5 mM CaCl_2_ and the turgid weight (TW) was recorded. Total dry weight (DW) was recorded after drying these leaves at 55°C for 48 hours. Leaf RWC was calculated as (FW−DW)/(TW−DW)x100.

### Soil volumetric water content (VWC)

VWC was measured using the 5TM soil moisture and temperature sensor, combined with the ‘ProCheck’ interface reader (Decagon Devices, Pullman, WA, USA).

### Measurements of leaf area

The plant’s total leaf area was measured with a Li 3100 leaf area meter (Li-Cor area meter, model Li 3100, Lincoln, NE, USA).

### Statistical analyses

All assays were conducted with three or more biological replicates and analyzed using JMP software (SAS Institute, Cary, NC). Means comparison was conducted using ANOVA with post-hoc Tukey-Kramer HSD (For multiple comparisons) and Student’s t tests (For one comparison) (P<0.05).

### Gene annotation and accession numbers

*GA2ox1* to *GA2ox5* were named here according to Pattison et al. (2015) and *GA2ox7* according to Schrager-Lavelle et al. (2019) and all other *GA2oxs* were named by their accession numbers. *GA20ox1* to *GA20ox4* and *GA3ox1* and *GA3ox2* were named here according to Pattison et al. (2015), all other *GA20oxs* and *GA3oxs* were named by their accession numbers.

Sequence data from this article can be found in the Sol Genomics Network (https://solgenomics.net/) under the following accession numbers: *ACTIN*, Solyc11g005330; *GA2ox1*, Solyc05g053340; *GA2ox2*, Solyc07g056670; *GA2ox3*, Solyc01g079200; *GA2ox4*, Solyc07g061720; *GA2ox5*, Solyc07g061730; *GA2ox7*, Solyc02g080120; Other *GA2oxs*: Solyc01g058040; Solyc02g070430; Solyc08g016660; Solyc10g007570; Solyc01g058030; *GA20ox1*, Solyc03g006880; *GA20ox2*, Solyc06g035530; *GA20ox3*, Solyc11g072310; *GA20ox4*, Solyc01g093980; Other *GA20oxs:* Solyc06g050110; Solyc09g009110; Solyc10g046820; Solyc11g013360; *GA3ox1*, Solyc06g066820 ;*GA3ox2*, Solyc03g119910; Other *GA3oxs*: Solyc01g058250; Solyc05g052740; Solyc00g007180; Solyc01g067620; *TINY1*, Solyc06g066540; Other TINYs: Solyc08g066660; Solyc12g044390; Solyc01g090560; Solyc12g008350; Solyc03g120840.

## SUPPLEMENTAL DATA

**Supplemental Figure 1**. Vitality of the isolated guard cells.

**Supplemental Figure 2**. *GA2ox7* exhibits the highest expression among all analyzed *GA2ox* genes in guard cells.

**Supplemental Figure 3**. Drought regulation of *GA3oxs* and *GA20oxs* expression in guard cells.

**Supplemental Figure 4**. Drought regulation of *GA2ox, GA3ox* and *GA20ox* in leaves.

**Supplemental Figure 5**. ABA regulation of GA metabolism genes

**Supplemental Figure 6**. Putative DREB-responsive elements (DRE) in the *GA2ox7* promoter.

**Supplemental Figure 7**. Sequence analyses of *tiny1* CRISPR-Cas9 mutants.

**Supplemental Figure 8**. Loss of *GA2ox7* affects stem elongation but not leaf growth.

**Supplemental Figure 9**. Loss of *GA2ox7* did not affect transpiration rate and stomatal aperture in well-watered plants.

**Supplemental Figure 10**. Loss of *GA2ox7* inhibited stomatal closure in response to mild soil dehydration.

**Supplemental Figure 11**. Sequence analyses of *ga20ox1 and ga20ox2* CRISPR-Cas9 mutants.

**Supplemental Figure 12**. Loss of *GA20ox1* and *GA20ox2* suppress growth but had no effect on basal stomatal aperture.

**Supplemental Figure 13**. The double mutant *ga20ox1/ga20ox2*.

**Supplemental Figure 14**. The double mutant *ga20ox1*/*ga2ox7* has long epicotyl, similar to *ga2ox7* and small leaves, similar to *ga20ox1*.

**Supplemental Table 1**. sgRNAs used in this study.

**Supplemental Table 2**. Primers used in this study.

**Supplemental Table 3**. UPLC-MRM parameters used for measuring gibberellins by UPLC-QQQ-MS/MS.

**Dataset 1**.GA analyses in leaves and guard cells.

## AUTHOR CONTRIBUTIONS

H.S, Y.E. and D.W. designed the research plan; H.S., H.C., H.V., N.I-E, S.B. and Z.A. performed the research; A.A and D.T contributed analytic tools; H.S., H.V., A.A. and D.T. analyzed data; H.S., Y.E and D.W. wrote the paper.

## ACKNOWLEDGEMENTS

This research was supported by the Israel Science Foundation (617/20) to D.W., and by the European Regional Development Fund Project “Centre for Experimental Plant Biology” (No. CZ.02.1.01/0.0/0.0/16_019/0000738) and the Czech Science Foundation (Grantová agentura České republiky, grant nr. 18-10349S) to D.T. We thank Prof. Zach Adam for his valuable suggestions. We also thank Dr. Sayantan Panda, Or Ram and Magdalena Vlckova for their technical assistance.

